# Two Interacting Neural Processes Support Speech Planning during Naturalistic Conversation

**DOI:** 10.64898/2026.05.06.723165

**Authors:** Hiroyoshi Yamasaki, Philippe Blache, Daniele Schön

## Abstract

Conversation requires speakers to plan their responses while still listening to their partner, yet the neural dynamics supporting this overlap remain unclear. In particular, competing accounts differ on whether behaviourally relevant preparation emerges early during comprehension or only close to speech onset. Here, we recorded EEG from pairs of participants engaged in natural, face-to-face conversation and tested whether neural activity during listening predicts both when speakers begin their response (latency) and how long they speak (duration). Using event-related potentials, oscillatory analyses, and multivariate decoding, we show that pre-speech neural activity carries robust information about upcoming behaviour more than one second before articulation. The strongest and earliest effects tracked response duration, with sustained ERP components and alpha/beta power modulations predicting how long participants would speak, even after controlling for behavioural variables. In contrast, neural predictors of response latency were more temporally restricted and partly aligned with partner turn boundaries. Temporal generalisation analyses further revealed stable neural patterns linking early and late stages of preparation. Together, these findings indicate that conversational planning unfolds during listening and that early neural activity reflects the maintained specification of response extent, followed by later processes related to commitment and initiation. This supports a temporally structured account of turn taking in which preparation involves both early maintenance and late motor engagement.

## Introduction

Conversation is characterised by a surprisingly efficient interpersonal coordination in which interlocutors switch turns with small latencies of only a few hundred milliseconds (Stivers et al., 2009). This speed is striking because speech production is thought to require substantially longer, often estimated at 600 ms or more from conceptual preparation to articulation (Indefrey and Levelt, 2004). The mismatch between rapid turn exchange and slow production bottlenecks implies that, in natural dialogue, speakers must begin preparing their own utterances while still listening to their partner (Levinson and Torreira, 2015). This observation has been a key motivation for influential accounts of dialogue that emphasise tight coupling between comprehension and production (Pickering and Garrod, 2013), but it also raises a more specific question: When, during ongoing comprehension does neural activity begin to reflect the timing and extent of the upcoming response, and how does this activity evolve relatively to turn boundary?

A central debate in the turn-taking literature concerns the timing and control architecture of response planning (see Fig. 1). The *Early Planning Hypothesis* proposes that listeners begin preparing their response as soon as sufficient information becomes available, potentially well before the partner has finished speaking (e.g., Levinson and Torreira, 2015). In contrast, the *Late Planning Hypothesis* emphasizes constraints on speech-motor initiation, suggesting that although some aspects of conceptual preparation may occur during listening, full articulatory planning is deferred until close to the end of the partner’s turn in order to minimise interference between comprehension and production (e.g., Sjerps and Meyer, 2015). Thus, the two views differ not in whether preparation occurs during listening, but in *when* it occurs. Under strong early planning accounts, neural activity predictive of the upcoming response should be observable during the listening interval, potentially more than a second before speech onset. Under late initiation accounts, predictive neural activity should be concentrated closer to articulation, reflecting the strategic timing of motor preparation (Bögels et al. 2015a).

**Figure 1.**
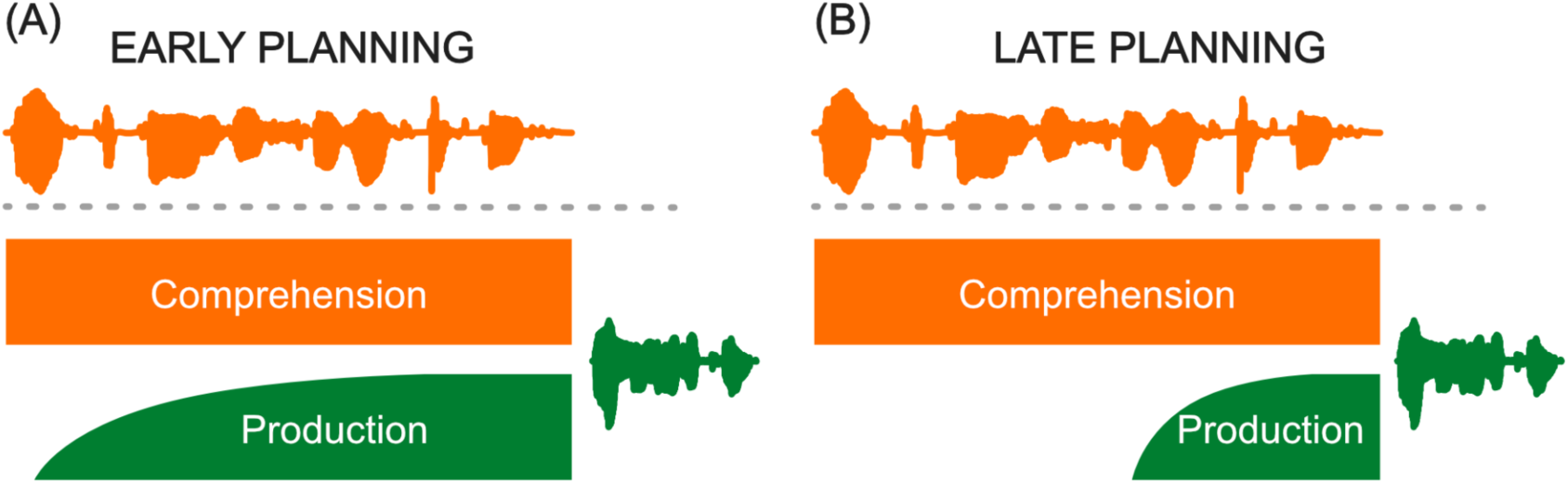
Early vs. Late Planning Hypothesis. (A) *Early Planning Hypothesis*. The speaker starts planning speech production as soon as enough information is available (B) *Late Planning Hypothesis*. The speaker defers planning speech production until near the end to avoid overlap

Behavioural evidence has supported both viewpoints. Some studies suggest that upcoming speakers can prepare response content early but wait for turn-end cues to initiate it (for example, Corps et al., 2018; Barthel et al., 2016, 2017), whereas others highlight situations in which response preparation is delayed (for example, Sjerps and Meyer, 2015; Boiteau et al., 2014). Neuroimaging and electrophysiology studies using indirect proxies of conversational behaviour tend to support early preparation, reporting preparatory alpha and beta desynchronisation and readiness-like slow potentials well before speech onset (for example, Berthault et al., 2026; Magyari et al., 2014; Piai et al., 2015). However, much of this evidence comes from controlled or semi-scripted tasks that differ from spontaneous conversation in both structure and demands. It therefore remains unclear to what extent these neural signatures track turn-taking behaviour in genuinely natural interaction.

A further limitation is that many neural studies of dialogue either use simulated interaction, where participants listen to recordings rather than engage in real-time exchange (for example, Stephens et al., 2010; Kuhlen et al., 2012), or rely on paradigms in which participants observe conversations, such as the overhearer approach (for example, Egorova et al., 2014, 2016; Bögels et al., 2015b; Bašnáková et al., 2014; Van Ackeren et al., 2012; Gisladottir et al., 2018). While these designs are valuable for isolating comprehension-related processes, they only partially engage the core computational challenge of turn taking, namely coordinating one’s own production with an unfolding partner turn. A small number of studies have examined neural activity in natural conversation, providing important insights into global dynamics (for example, Pérez et al., 2017; Bellegarda et al., 2025), but they did not focus on the fine-grained temporal structure of turn exchange. Even when temporal dynamics are addressed, analyses target different representational levels, such as word embeddings, rather than turn-level planning variables (for example, Zada et al., 2024). As a result, it remains unclear how pre-speech neural activity during listening relates to the timing and structure of the next spoken response in genuine interaction.

To date, the clearest electrophysiological evidence directly tied to turn-taking preparation comes from work by Bögels and colleagues. Using controlled quiz-style exchanges, they reported a slow posterior positivity and alpha desynchronisation beginning up to about 2 seconds before speech onset (Bögels et al. 2015a, 2018; Bögels, 2020). These findings are consistent with early preparation. However, because these paradigms primarily emphasised processing the incoming question rather than predicting and controlling the timing and extent of a spontaneous response, it remains unclear how well they capture turn-taking in more natural settings. Because behavioural dynamics are fundamental to linking neural activity with communicative function (Krakauer et al., 2017), progress on the early versus late planning debate requires neural analyses that are both ecologically grounded and behaviour-linked at the level of individual turns.

Here we address this gap in the literature by recording EEG while pairs of participants engaged in fully naturalistic, face-to-face conversation under minimal experimental constraints. We operationalised turn-taking behaviour using two complementary measures that place constraints on competing timing accounts: response latency and response duration. Response latency, defined as the gap between the partner’s offset and the speaker’s onset, is the most widely used behavioural index in the turn-taking literature and has been central to characterising the temporal precision of conversational exchange (see Levinson and Torreira, 2015 for a review). Response duration, defined as the length of the upcoming contribution, has received comparatively little attention (although see Torreira et al. 2015), despite potentially being relevant to turn-taking behaviour. Although these variables are coupled in natural dialogue and influenced by partner behaviour, they are not reducible to one another. Latency primarily reflects initiation control at the turn boundary, whereas duration reflects the scale and persistence of the forthcoming response. Considering both measures therefore provides a more complete characterisation of turn-taking behaviour and offers leverage to test whether early neural activity carries information about upcoming behaviour beyond conversational context.

Our central aim was therefore to determine whether neural signals recorded while a participant listens to their partner predict when they will begin speaking and how long they will speak, and to characterise the time course of these predictive signals. We aligned EEG to the onset of each utterance and analysed pre-speech activity using event-related potentials. We also looked at alpha and beta band powers because of their importance in turn taking according to the previous works (Bögels et al. 2015a, 2018; Bögels, 2020) as well as their relevance for motor planning (Démas et al. 2020). These analyses were complemented by trial-level modelling that controls key conversational covariates, including partner turn characteristics and the corresponding behavioural variables. By linking neural activity in defined pre-speech windows to continuous measures of latency and duration, we test whether behaviour-relevant preparation emerges early during listening, whether additional predictive dynamics appear closer to speech onset, and how these patterns jointly constrain timing-based accounts of conversational planning.

## Results

### Conversational timing is mutually adaptive

We first characterised the behavioural structure of turn taking in our dataset, because these dependencies determine what must be controlled when testing neural predictors. Across 12,939 turns, we defined response latency as the time from the partner’s IPU offset to the participant’s IPU onset, with negative values indicating temporal overlap, and response duration as the length of the participant’s upcoming IPU (see Fig. 2). For the sake of brevity, in the following the participants whose EEG data is being analysed will be referred to as *self*-participants and their partner as *other*-participants.

**Figure 2.**
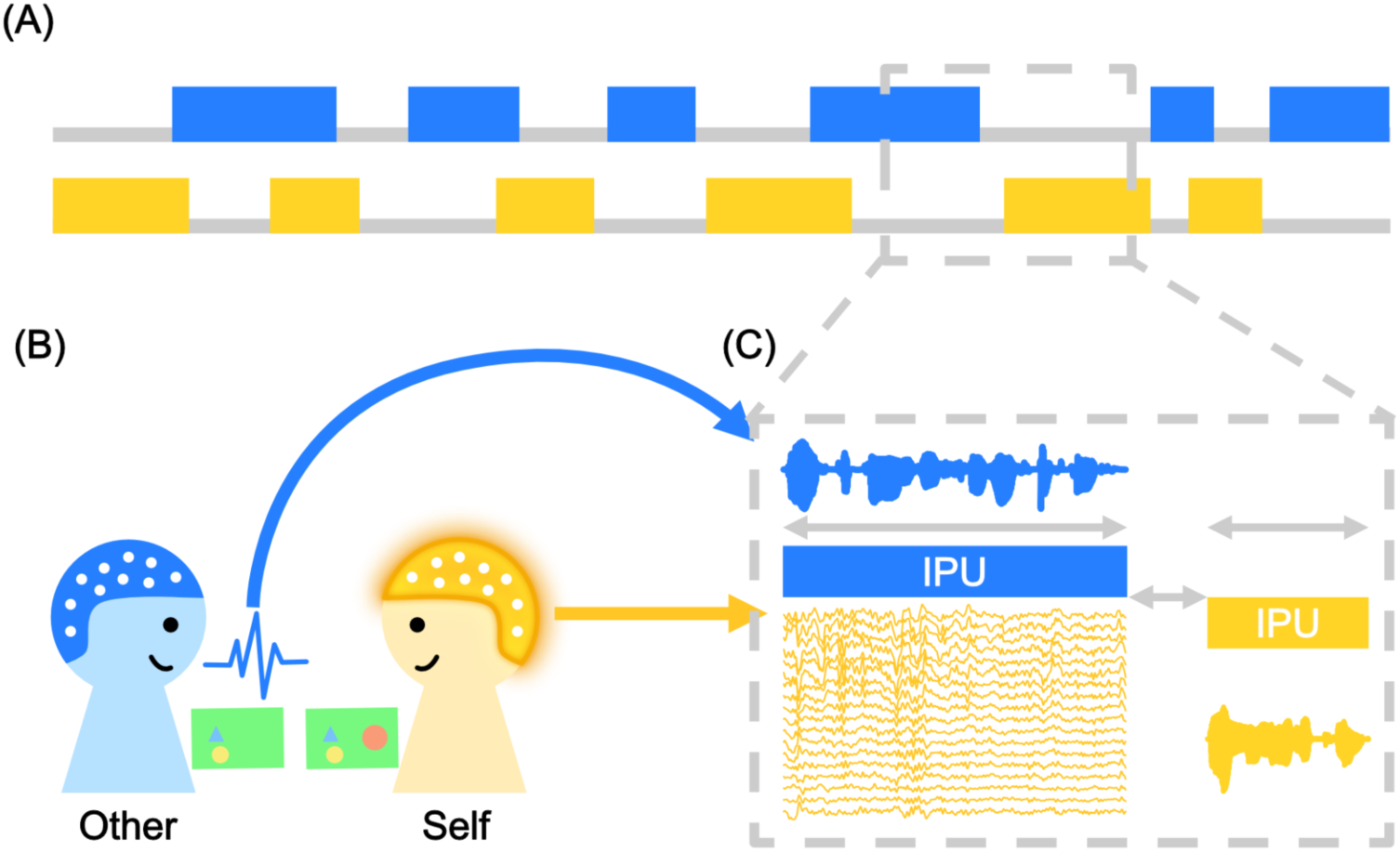
Experimental setup and analysis design. (A) The conversation is segmented into *inter-pausal units* (IPUs), defined as stretches of continuous speech produced by a single speaker and bounded by silent pauses of at least 200 ms. IPUs serve as the basic units for defining turn transitions and behavioural measures (response latency and duration). (B) For each analysis, participants are designated as either *self* (yellow) or *other* (blue). *Self* refers to the participant whose EEG data are currently being analysed, whereas *other* refers to their conversational partner. Because analyses are performed symmetrically for both members of the dyad, each participant serves as *self* during their own listening periods and as *other* during their partner’s. (C) EEG analyses focus exclusively on the listening interval preceding the self-participant’s speech onset, thereby isolating preparatory neural activity before articulation.

Response duration showed substantial variability and was well described by a log-normal shaped distribution (see Fig. 3). This wide spread indicates that speakers produced both brief and extended contributions, providing meaningful variation in how long they continued once they took the floor. Response latencies were distributed around a clear central tendency, with a mode around 280 ms, consistent with previous corpus estimates of floor transition timing (Levinson and Torreira, 2015). Together, these distributions confirm that our conversations display the characteristic fast transitions of natural dialogue, while also containing meaningful variation in both how long speakers continue and when they initiate their turns.

**Figure 3.**
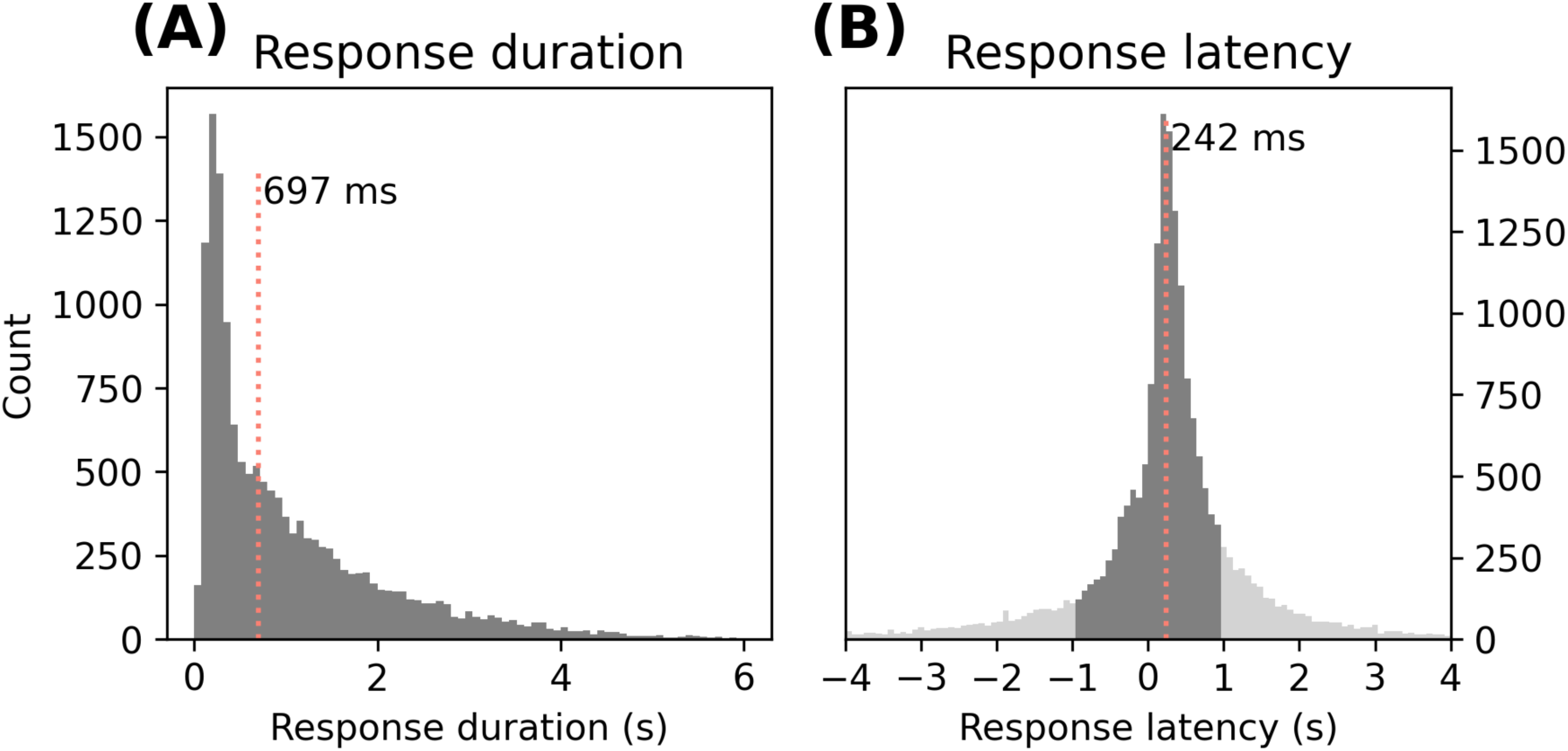
Distributions of behavioural measures across all participants. (A) Response duration, defined as the length of the speaker’s IPU. Only trials included in subsequent EEG analyses are shown (dark gray), corresponding to turn-taking events meeting the latency inclusion criteria. Vertical dashed lines mark the median used for condition splits. (B) Response latency, defined as the time from the offset of the partner’s IPU to the onset of the speaker’s IPU. Negative values indicate temporal overlap between speakers. Dark gray bars indicate trials included in subsequent EEG analyses, whereas light gray bars indicate excluded trials.

Latency and duration were not independent. Mixed-effects modelling revealed robust couplings that reflect mutual adaptation between interlocutors. Longer responses tended to follow longer latencies (self-duration positively related to latency, β = 0.12 ± 0.0072, t = 17.23). In addition, turn lengths covaried across speakers in a compensatory manner, such that longer turns by one partner tended to elicit shorter responses from the other (self-duration negatively related to other-duration, β = −0.13 ± 0.0072, t = −17.23). Finally, longer partner turns were followed by faster responses (latency negatively related to other-duration, β = −0.028 ± 0.0073, t = −3.87) which replicates previous results (Bögels and Torreira 2015).

Because latency and duration are both influenced by the immediately preceding partner turn, and because latency and duration also covary with each other, neural effects on either behavioural outcome can be confounded if these behavioural dependencies are ignored. We therefore include partner duration (other-duration) as a core covariate in all neural linear mixed effect models, along with the reciprocal behavioural variable (latency when predicting duration, and duration when predicting latency). This covariate structure allows us to test whether pre-speech neural activity during listening explains additional variance in upcoming behaviour beyond what is already predictable from conversational context and the intrinsic coupling between when speakers start and how long they speak.

### Pre-speech ERPs during listening show sustained activity well before speech onset

To characterise preparatory activity preceding each response, we analysed ERPs time-locked to the onset of the participant’s own IPU (speech onset at 0 s). As illustrated in Fig. 4, response-locked ERPs appear to vary with both response duration and response latency. These descriptive patterns motivated a subsequent data-driven clustering approach to identify the spatiotemporal regions of interest used in later analyses. Specifically, we applied cluster-based permutation tests across the full epoch and all electrodes to isolate time windows and regions showing reliable modulation by latency and duration, splitting trials according to median response duration (long versus short) and response latency (slow versus fast). For the duration comparison (long minus short), cluster-based testing revealed four significant spatio-temporal clusters (cluster-corrected p < 0.01, Maris and Oostenveld 2007). Two large clusters extended from roughly −1200 ms to −300 ms before speech onset and consisted of a frontal negativity together with a posterior positivity, resembling a slow-wave pattern reported previously in more controlled response-planning paradigms (Berthault et al., 2026, Bögels et al., 2015a, 2018; Bögels, 2020). Two additional clusters occurred close to speech onset (approximately −100 to 0 ms, p = 0.006) and showed reversed polarities (see Fig. 5 & Supplementary Fig. S1, S4).

**Figure 4.**
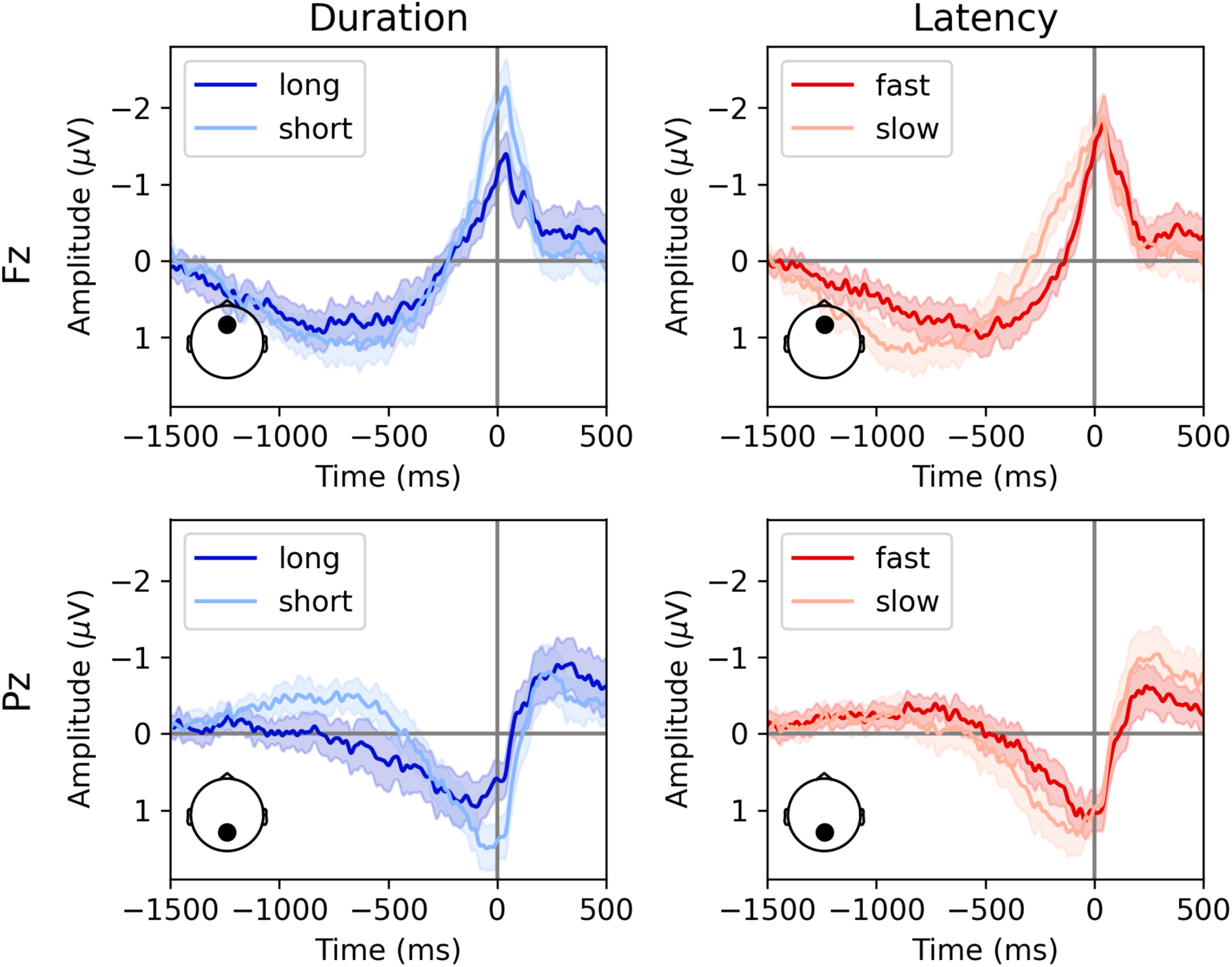
Event-related potentials (ERPs) by condition in Fz (top) and Pz (bottom) electrodes. (Left) ERPs time-locked to response onset for long versus short upcoming IPU durations (inter-pausal units). (Right) ERPs time-locked to response onset for fast versus slow response latencies. Shaded areas indicate the 95% confidence interval across participants (N = 32).

**Figure 5.**
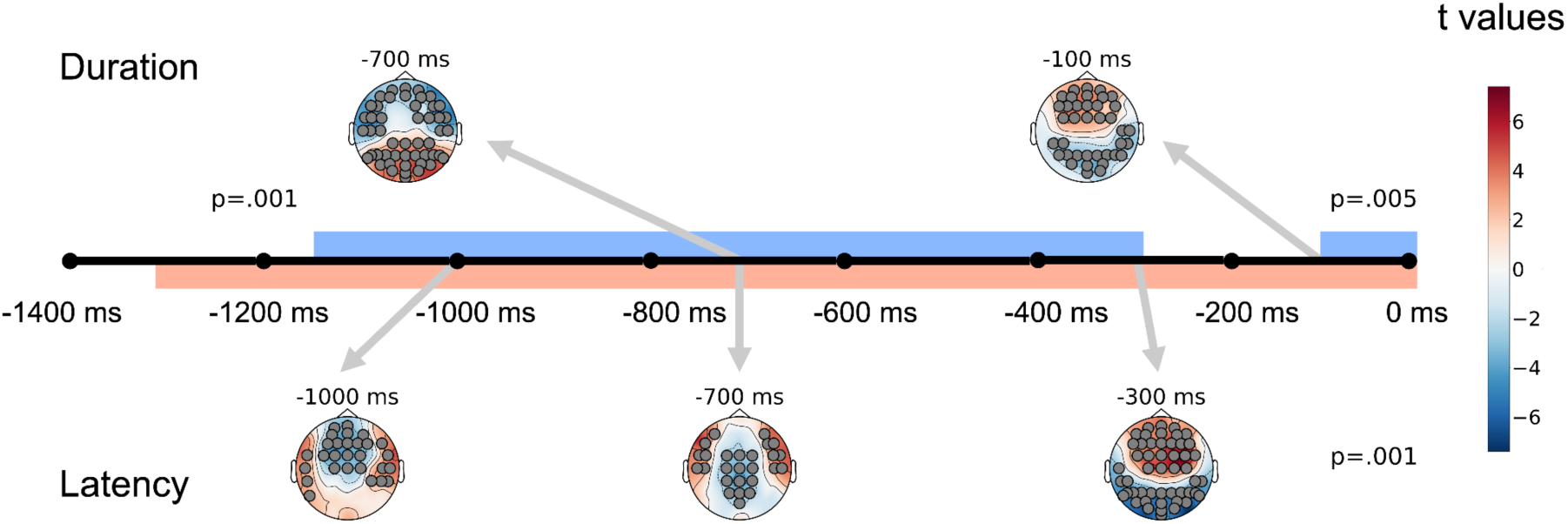
Summary of ERP effects of response duration and response latency. Topographies of *t*-values for the duration (top) and latency (bottom) contrasts, time-locked to speech onset. Warm and cool colours indicate relative positivity or negativity for the first condition in each contrast (long - short; fast - slow). Black circles mark electrodes belonging to significant spatio-temporal clusters identified by cluster-based permutation tests (*p* < 0.05).

For the latency comparison (fast minus slow), cluster-based testing revealed two broad spatio-temporal clusters (cluster-corrected p < 0.001), spanning approximately −1300 ms to speech onset. One cluster showed an anterior negativity, and the other a lateral-posterior positivity (see Fig. 5 & Supplementary Fig. S1, S5). The anterior effect appeared closely aligned with the timing of the partner’s speech offset, suggesting a potential contribution from auditory or turn-boundary related responses, whereas the posterior effect showed a temporal profile consistent with sustained preparatory activity leading up to the participant’s own onset.

To further assess whether latency-related ERP effects reflected offset-locked auditory responses rather than initiation-related preparation, we examined the temporal alignment between the partner’s speech offset and the ERP inflection points in anterior and posterior regions (see Supplementary Fig. S6). In anterior electrodes, the ERP inflection closely tracked the distribution of partner offset timing, whereas no such alignment was observed in posterior sites. This dissociation suggests that the anterior latency effect may partly reflect offset-evoked auditory activity, while the posterior effect is less likely to be driven by partner-offset timing.

We treat these condition-split ERP contrasts as descriptive anchors to identify spatio-temporal regions of interest (ROI) and pre-speech time-window which are then carried forward into trial-level modelling.

### ERP amplitude predicts upcoming turn duration and latency beyond conversational covariates

To test whether ERP activity during listening predicts upcoming behaviour beyond conversational context, we quantified, for each trial, mean ERP amplitudes within the regions and time windows identified by the cluster-based analyses, and entered these into linear mixed-effects models (see Table 1). ERP predictors were defined as mean amplitude within two pre-speech windows (TW1: −1400 to −800 ms; TW2: −800 to 0 ms) and within anterior and posterior regions of interest. Models included behavioural covariates capturing conversational context, namely partner turn duration (other-duration) and the reciprocal behavioural variable (response latency when predicting duration, and response duration when predicting latency). Neural effects were evaluated by likelihood-ratio tests comparing models with and without the ERP term.

**Table 1.** Summary of Mixed Effects Model Results. Each row reports the effect of a neural predictor extracted during the listening interval before self speech onset. Neural predictors comprised ERP amplitude, alpha-band activity, and beta-band activity, each summarised within a region of interest (ROI: anterior or posterior) and a pre-speech time window (TW1: −1400 to −800 ms; TW2: −800 to 0 ms). Separate models were fitted for the two behavioural outcomes: upcoming response duration and upcoming response latency. β indicates the standardised fixed-effect coefficient for the neural predictor, SE its standard error, and χ² the likelihood-ratio test statistic obtained by comparing the full model including the neural predictor with a corresponding null model without that predictor. *FDR p* denotes the false-discovery-rate-corrected p-value across all neural tests within each outcome analysis. The *sig.* column indicates whether the neural effect survived FDR correction at α = 0.05. All models included participant as a random intercept and controlled for partner duration (*other-duration*) and the reciprocal behavioural variable (latency when predicting duration; duration when predicting latency)

| Predictor | TW | ROI | $\beta$ | SE | $\chi^2$ | FDR p | Sig. |
| --- | --- | --- | --- | --- | --- | --- | --- |
| **Duration** |  |  |  |  |  |  |  |
| ERP | TW1 | Anterior | -0.009 | 0.008 | 1.19 | 0.275 | nan |
| | TW1 | Posterior | 0.047 | 0.008 | 30.86 | $1.11 \times 10^{-7}$ | *** |
|  | TW2 | Anterior | -0.021 | 0.009 | 6.23 | 0.019 | * |
| | TW2 | Posterior | 0.056 | 0.009 | 42.13 | $1.03 \times 10^{-9}$ | *** |
| Alpha | TW1 | Anterior | -0.019 | 0.010 | 4.05 | 0.059 | nan |
|  | TW1 | Posterior | -0.026 | 0.010 | 7.52 | 0.010 | * |
|  | TW2 | Anterior | -0.019 | 0.010 | 3.61 | 0.064 | nan |
| | TW2 | Posterior | -0.038 | 0.010 | 13.77 | $4.96 \times 10^{-4}$ | *** |
| Beta | TW1 | Anterior | -0.021 | 0.011 | 3.59 | 0.064 | nan |
|  | TW1 | Posterior | -0.036 | 0.011 | 10.73 | 0.002 | ** |
| | TW2 | Anterior | -0.056 | 0.011 | 23.59 | $3.57 \times 10^{-6}$ | *** |
| | TW2 | Posterior | -0.073 | 0.012 | 35.46 | $1.56 \times 10^{-8}$ | *** |
| **Latency** |  |  |  |  |  |  |  |
| ERP | TW1 | Anterior | 0.065 | 0.009 | 56.09 | $4.16 \times 10^{-13}$ | *** |
|  | TW1 | Posterior | -0.008 | 0.009 | 0.95 | 0.361 | nan |
| | TW2 | Anterior | -0.043 | 0.009 | 24.37 | $3.18 \times 10^{-6}$ | *** |
| | TW2 | Posterior | 0.067 | 0.009 | 56.96 | $4.16 \times 10^{-13}$ | *** |
| Alpha | TW1 | Anterior | 0.024 | 0.010 | 5.77 | 0.028 | * |
| | TW1 | Posterior | 0.039 | 0.010 | 16.01 | $1.90 \times 10^{-4}$ | *** |
|  | TW2 | Anterior | 0.004 | 0.010 | 0.13 | 0.721 | nan |
|  | TW2 | Posterior | 0.034 | 0.010 | 10.89 | 0.002 | ** |
| Beta | TW1 | Anterior | 0.017 | 0.011 | 2.20 | 0.165 | nan |
|  | TW1 | Posterior | 0.021 | 0.011 | 3.34 | 0.102 | nan |
|  | TW2 | Anterior | -0.018 | 0.012 | 2.52 | 0.150 | nan |
|  | TW2 | Posterior | -0.032 | 0.012 | 6.58 | 0.021 | * |

Even after controlling for the aforementioned behavioural variables, ERP amplitude significantly predicted upcoming response duration, with the most consistent effects in posterior electrodes across both time windows. Anterior effects did not survive correction (see Table 1). These findings indicate that pre-speech ERP activity during listening carries information about how long the participant will speak, beyond what is explained by partner behaviour and the intrinsic latency-duration coupling.

ERP amplitude also predicted upcoming response latency. Anterior effects were present in both time windows, and posterior effects were strongest in the later window. Notably, duration-related prediction was already evident in the earlier window, whereas latency-related effects were more prominent closer to speech onset (see Table 1).

Overall, these covariate-controlled models show that ERPs during the listening interval contain behaviour-predictive information about both response extent and initiation timing.

### Alpha and beta power show pre-speech modulations during listening

We next examined oscillatory activity during the same pre-speech listening interval using time-frequency analyses focused on the alpha (8-12 Hz) and beta (13-30 Hz) bands. Previous studies have shown that power in these frequency bands is associated with turn-taking behaviour (Bögels et al., 2015a, 2018; Bögels, 2020). As for the ERP analyses, we summarised effects relative to the participant’s speech onset (0 s) and assessed condition differences using cluster-based spatio-temporal permutation testing across sensors.

For the duration comparison (long minus short), alpha-band power showed a sustained pre-speech modulation. A significant alpha cluster (cluster-corrected p < 0.001) emerged at approximately -1.2 s before speech onset and extended towards articulation, with strongest expression over frontal and parietal electrodes (see Fig. 6 & Supplementary Fig. S2). The direction of the effect reflected reduced alpha power for longer upcoming turns relative to shorter turns.

**Figure 6.**
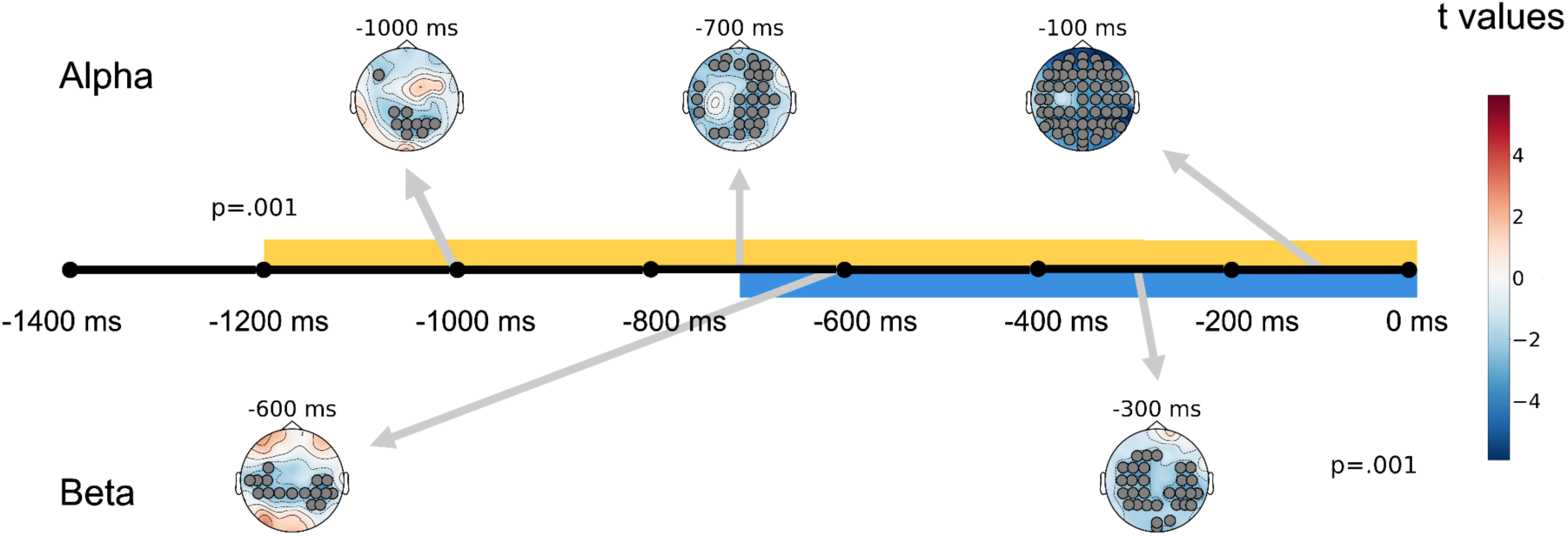
Alpha/beta power modulation by response duration. Topographies of *t*-values for the duration contrast in alpha (top) and beta (bottom) bands, time-locked to speech onset. Warm and cool colours indicate relative positivity or negativity (long - short). Black circles mark electrodes belonging to significant spatio-temporal clusters identified by cluster-based permutation tests (*p* < 0.05).

Beta-band power also differed as a function of upcoming duration, with a later time course (see Fig. 6 & Supplementary Fig. S3). A significant beta cluster (cluster-corrected p < 0.001) was concentrated over central electrodes and peaked within the final −0.7 to 0 s before speech onset. As in the alpha band, beta power was reduced for longer upcoming turns.

For the latency comparison (slow minus fast), no significant clusters were observed in either the alpha or beta bands in the condition-split contrasts. These time-frequency results therefore show robust pre-speech alpha and beta modulations associated with the duration of the upcoming response during the listening interval, while corresponding condition-level differences for latency were not detected with this approach.

### Alpha and beta power predict upcoming behaviour beyond conversational covariates

We next asked whether oscillatory power during listening predicted upcoming behaviour beyond conversational context using trial-level linear mixed-effects models (see Table 1). Alpha- and beta-band predictors were defined as mean power within the same two pre-speech windows as for ERP analyses (TW1: -1400 to -800 ms; TW2: -800 to 0 ms) and within anterior and posterior regions of interest. Models included participants as a random intercept and the same behavioural covariates used in the ERP analyses. Neural effects were assessed by likelihood-ratio tests comparing models with and without the oscillatory predictor.

In the alpha band, posterior power reliably predicted both response duration and response latency. Reduced posterior alpha during listening was associated with longer upcoming responses in both time windows. Posterior alpha also predicted response latency, such that lower alpha was associated with slower turn initiation. Anterior alpha effects were weaker and less consistent across windows.

In the beta band, effects were more selective. Reduced beta power robustly predicted longer upcoming response duration across regions and time windows. In contrast, beta power did not significantly predict response latency in any window or ROI.

Taken together, these results indicate a dissociation between alpha and beta dynamics. Alpha activity during listening relates to both when speakers begin and how long they speak, whereas beta activity primarily tracks variation in response duration. Importantly, these relationships hold after controlling for partner behaviour and the intrinsic coupling between latency and duration (Table 1).

### Decoding reveals temporally stable neural patterns predictive of duration and latency

To characterise the temporal dynamics of behaviour-related neural information beyond univariate summaries, we performed temporal generalisation decoding on the EEG epochs (King and Dehaene, 2014). This approach allows us to assess whether neural representations are transient (rapidly changing over time) or temporally stable (maintained across time). We trained linear classifiers at each time point to discriminate short versus long upcoming responses and fast versus slow upcoming responses, and then tested each classifier across all time points to obtain temporal generalisation matrices. Performance was quantified using area under the ROC curve (AUC), and statistical significance was assessed using cluster-based permutation tests against chance performance (AUC = 0.5).

Decoding accuracy for response duration exceeded chance along the diagonal from approximately −1300 ms before speech onset and increased towards articulation, peaking around the onset period (see Fig. 7). Temporal generalisation matrices revealed a large, nearly square region of above-chance decoding spanning roughly −1300 ms to 0 ms, indicating that the neural patterns predictive of upcoming duration were temporally stable across much of the pre-speech interval (King and Dehaene, 2014).

**Figure 7.**
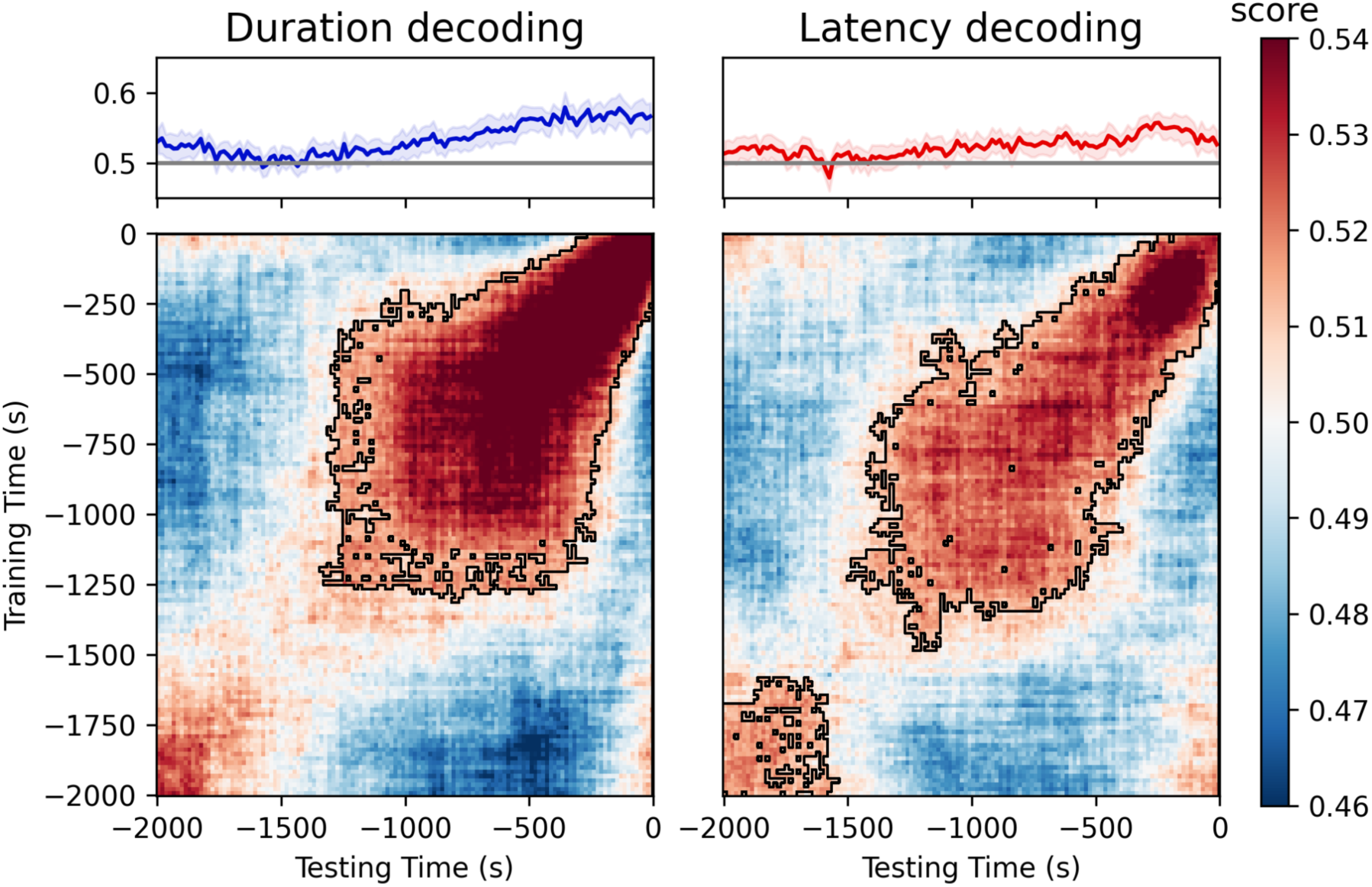
Temporal generalisation decoding of response duration and latency. (A) Diagonal decoding accuracy (area under the ROC curve, AUC) for classifying short vs. long responses (left) and fast vs. slow responses (right). Shaded areas indicate 95% confidence interval across participants; The black outlines mark the statistically significant clusters (*p* < 0.05). (B) Temporal generalisation matrices showing decoding performance across training (y-axis) and testing (x-axis) times relative to speech onset (0 s). Red regions indicate above-chance classification and black contours denote significant clusters. Both duration and latency could be decoded from neural activity beginning ∼1.3 s before speech onset, with large, temporally stable patterns extending up to articulation.

Decoding accuracy for response latency also rose above chance for much of the pre-speech period, with the highest values occurring close to speech onset. Temporal generalisation was sustained from roughly −1200 ms to 0 ms, again consistent with a stable neural pattern linking earlier and later stages of response preparation. In addition, a smaller early cluster was observed between −2000 and −1500 ms, although interpretation of effects in this extreme early window is constrained by its distance from the turn boundary and by the broader variability in conversational context.

Overall, these decoding results show that information predictive of both how long speakers will talk and when they will begin speaking is present in pre-speech EEG activity and is maintained in a temporally stable format across a substantial portion of the listening interval. This representational stability complements the univariate ERP and oscillatory findings while avoiding strong assumptions about the specific neural mechanisms generating the predictive patterns.

### Distinct functional contributions of ERP and alpha to response duration

The preceding analyses revealed two key patterns. First, decoding suggested temporally stable pre-speech neural representations. Second, ERP amplitude and alpha power both predicted upcoming behaviour, but in partially opposing directions: ERP amplitude was positively related to response duration, whereas alpha power showed a negative relationship with duration. We therefore asked whether ERP amplitude and alpha power were themselves systematically related at the trial level.

To test this, we examined whether posterior alpha power during listening could be predicted from posterior ERP amplitude using a mixed-effects model controlling for behavioural covariates. Posterior alpha power was positively associated with ERP amplitude (β = 0.039, SE = 0.0077, χ²(1) = 26.1, p < 10⁻^6^), indicating that trials with larger ERP amplitudes tended to show higher alpha power during the same pre-speech interval.

Given the opposing behavioural associations of ERP and alpha, we next asked whether alpha explained variance in response duration beyond ERP amplitude. Adding alpha to a model already containing ERP and behavioural covariates significantly improved model fit (β = −0.029, SE = 0.0096, χ²(1) = 8.99, p = 0.0027). Importantly, the alpha coefficient was negative when ERP was included, indicating that higher alpha predicted shorter upcoming utterances after accounting for ERP amplitude.

Together, these results establish three empirical observations: ERP amplitude and alpha power covary positively; ERP amplitude predicts longer upcoming responses; and alpha power predicts shorter upcoming responses when ERP amplitude is held constant. These relationships constrain how slow ERP activity and alpha oscillations jointly relate to upcoming speech behaviour.

## Discussion

In this study, we show that neural activity during listening predicts properties of the next speaking turn in spontaneous, face-to-face conversation. Most robustly, pre-speech activity more than one second before articulation scales with the duration of the upcoming contribution, indicating that the extent of a response is represented while the partner is still speaking. These duration-related signals were evident in sustained slow ERPs, alpha and beta dynamics, and temporally stable multivariate patterns, and they remained significant after controlling for partner behaviour and latency-duration coupling. In contrast, neural predictors of response latency were more temporally constrained and partly aligned with partner turn boundaries. Together, these findings indicate that conversational planning unfolds during the listening interval and that early neural activity reflects the maintained specification of response extent, indexed here by response duration, rather than only the timing of speech initiation. Importantly, the present results do not allow us to conclude that these early signals are linguistic in nature in the sense of encoding specific lexical or semantic content. Instead, they index response-related parameters, as captured here by upcoming duration and latency, which may reflect the scale or complexity of the planned contribution.

### Early pre-speech activity reflects response extent

While the early vs late planning debate has primarily focused on the timing of linguistic content planning, the present study addresses a related but distinct question by examining when neural activity predicts response-related parameters such as duration and latency. The most robust result concerned response duration. A sustained slow ERP component beginning more than 1000 ms before speech onset scaled with how long participants were about to speak. This pattern resembles slow preparatory potentials reported in more controlled turn-taking and speech-planning paradigms, where pre-speech activity has been interpreted as reflecting anticipatory processes (Berthault et al., 2026; Bögels et al., 2015a, 2018; Bögels, 2020). However, in the present data the strongest early effect tracked not when participants began speaking, but how long they would continue. Because response duration covaries with partner turn duration (see behavioural analyses), sustained pre-speech activity could in principle reflect variation in listening duration. However, ERP effects remained significant after controlling for partner duration in mixed-effects models, indicating that they track properties of the upcoming response beyond the length of the preceding turn.

One natural point of reference is the readiness potential and related slow potentials linked to preparation for action (Schurger et al., 2012; Shibasaki and Hallett, 2006). Within *accumulator models*, such slow activity reflects evidence accumulation toward a decision threshold, with movement initiation occurring once activity reaches a bound (Schurger et al., 2012; Anders et al., 2015). In paradigms where the behavioural question concerns whether and when a response is initiated, and where the motor output is relatively stereotyped (e.g., a button press or a short, fixed utterance), this framework provides a coherent account of pre-response build-up and its relationship to initiation timing.

However, accumulator models primarily explain action initiation. They do not typically predict that the magnitude of pre-response activity should scale with the *extent* of the upcoming action. In our data, the early slow ERP component scaled with how long participants would speak, even after controlling for latency. This dissociation suggests that the signal reflects more than the approach to an initiation threshold. Rather than indexing only the decision to start speaking, it appears to track the degree to which a forthcoming response is specified and sustained during listening.

These findings are more consistent with the idea that the early ERP component indexes maintained response activation during listening rather than solely reflecting the approach to an initiation/motor threshold. Three empirical features support this view: (i) the early onset of the effect, well before speech, (ii) its temporal stability across the pre-speech interval, and (iii) its scaling with response duration rather than only with initiation timing.

Frameworks that emphasise competition and dynamic binding of distributed feature sets provide one way to conceptualise these properties (Roelfsema, 2023). In such accounts, multiple task-relevant feature sets can be active in parallel while competing for large-scale coordination, but only one configuration at a time becomes sufficiently integrated to drive overt action. This architecture allows response representations to be activated and maintained during comprehension without triggering premature execution.

Applied to turn taking, longer responses typically involve maintaining a larger or more structured set of propositions, lexical items, or planned sequences. Maintaining a richer candidate response during the listening interval would therefore increase overall activation of that configuration, consistent with the larger sustained ERP deflection observed here. The temporal generalisation results, which revealed stable neural patterns predictive of duration across much of the pre-speech interval, further support the idea of maintained activation rather than a brief trigger process near the turn boundary. On this view, the early slow ERP activity may reflect the degree to which a forthcoming response is already assembled and sustained during listening.

### Oscillatory dynamics: maintenance and commitment

Alpha and beta oscillations showed complementary dynamics. Reduced alpha and beta power during the listening interval predicted longer upcoming turns, with the strongest desynchronization emerging in the final ∼800 ms before speech onset, with a central, mu-like topography consistent with sensorimotor involvement (Pfurtscheller and Lopes da Silva 1999). Although we cannot determine whether this reflects content-related complexity, general planning demands, or motor preparation, this pattern is broadly consistent with prior work reporting alpha decreases before responses in controlled conversational paradigms (Bögels et al., 2015a, 2018; Bögels, 2020).

Alpha activity has been linked to the stabilisation of task-relevant representations and the gating of interference (Klimesch, 2012; Jensen and Mazaheri, 2010). In the earlier portion of the listening interval, alpha modulation may therefore contribute to maintaining partially specified response representations, as indexed here by upcoming duration, while comprehension continues. As speech onset approaches, the stronger alpha and beta desynchronisation is consistent with increasing motor preparation and articulatory readiness, in line with evidence linking beta dynamics to planned speech and motor sequencing (Piai et al., 2015; Démas et al., 2020).

Importantly, ERP amplitude and alpha activities were related but not redundant. Slow ERP activity strongly predicted response duration, and alpha explained additional variance beyond ERP amplitude.

One interpretation consistent with these findings is that both measures partly reflect sustained response activation, while alpha additionally indexes the system transition toward execution. In this view, early pre-speech ERP activity primarily is consistent with the maintained specification of a candidate response during listening. Alpha power, although influenced by this maintained activation, additionally captures the system’s shift toward commitment and motor readiness as speech onset approaches.

This interpretation naturally accounts for two aspects of the data. First, alpha predicted response duration even when ERP amplitude was held constant, indicating that it carries information beyond slow ERP activity alone. Second, the strongest alpha and beta desynchronisation occurred close to speech onset, consistent with a role in late-stage motor preparation rather than earlier attentional orienting. This contrasts with prior work reporting stimulus-locked alpha/beta effects interpreted as attentional shifts (Bögels et al., 2015a, 2018; Bögels, 2020), whereas here the response-locked timing suggests that these oscillatory changes track processes leading up to articulation. Rather than replacing earlier maintenance processes, this late modulation appears to mark a shift from sustaining a planned response to committing it for execution.

### Latency effects and boundary timing

Neural predictors of response latency were also observed, particularly in the period close to speech onset. However, their interpretation requires caution. In anterior electrodes, ERP inflection points aligned closely with the timing of the partner’s speech offset, suggesting that part of the latency-related activity may reflect offset-locked auditory or turn boundary processing rather than purely initiation planning.

Although mixed-effects models showed that neural measures predicted latency beyond behavioural covariates, the absence of robust condition-level oscillatory clusters for latency and the temporal alignment with partner offset constrain strong claims about dedicated initiation signal during the earlier listening interval. Compared to the robust duration-related effects, latency predictors were more temporally restricted and more closely tied to boundary timing.

Taken together, these findings suggest that while aspects of initiation timing may be prepared during listening, the most robust early neural signature concerns the extent of the upcoming response rather than the precise moment of turn onset.

### Reconciling Early and Late Planning through maintenance and commitment

The present findings help clarify the longstanding debate between Early and Late Planning accounts of turn taking. Early planning theories propose that response preparation begins as soon as sufficient information becomes available, whereas late planning accounts emphasise strategic timing of motor initiation close to the turn boundary.

Our results suggest that these accounts do not need to be mutually exclusive, but instead describe distinct phases of a temporally structured process. Early activity during listening appears to reflect the activation and maintenance of response-related information, including the extent of the forthcoming contribution, as approximated here by response duration. Duration-related ERP effects emerged more than one second before speech onset and were sustained across the listening interval, indicating that aspects of the upcoming response are specified while the partner is still speaking. However, the present analyses do not allow us to determine whether these signals reflect linguistic content, general planning demands, or motor preparation. Within a competition account, the partner’s ongoing speech remains the dominant driver of perception, while a candidate response is partially activated in parallel. This interpretation also helps reconcile our findings with accounts proposing competition between comprehension and production processes (e.g. Sjerps and Meyer 2015). Early response-related activity may reflect partially activated representations that do not yet strongly compete with ongoing perception. Competition would then emerge more strongly at later stages, as response representations become increasingly committed to motor execution. This is also consistent with the temporal generalisation results, which indicate that response-related information can be maintained as a relatively stable neural pattern across much of the listening interval.

A complementary set of findings point to a later phase that is more closely related to commitment and execution. Evidence for Late Planning has often come from dual-task paradigms, where interference with concurrent tasks increases near the end of the partner’s turn and around speech onset (Barthel et al., 2016, 2017; Sjerps and Meyer, 2015; Boiteau et al., 2014). In our data, alpha and beta desynchronisation became stronger in the final ∼800 ms before speech onset, with a central, mu-like topography compatible with increasing motor involvement. This is consistent with a transition from maintaining a candidate response to preparing it for overt production.

On this view, Early and Late Planning accounts capture different functional components of conversational speech preparation. Early processes support the activation and maintenance of response-related representations, including the scale (duration) of the forthcoming contribution. Later processes reflect a shift toward execution, as attentional resources are reallocated and motor systems become increasingly engaged. Larger upcoming responses would therefore be expected to place greater demands both on maintained activation and on motor preparation, leading to stronger early ERP effects and more pronounced onset-proximal oscillatory changes.

Rather than choosing between Early and Late Planning, the present results suggest a temporally structured process in which maintenance and commitment unfold sequentially but partially overlap during natural conversation. In this framework, early preparation supports what and how much to say, while later processes govern when and how to implement the response. This interpretation is summarised schematically in Fig. 8.

**Figure 8.**
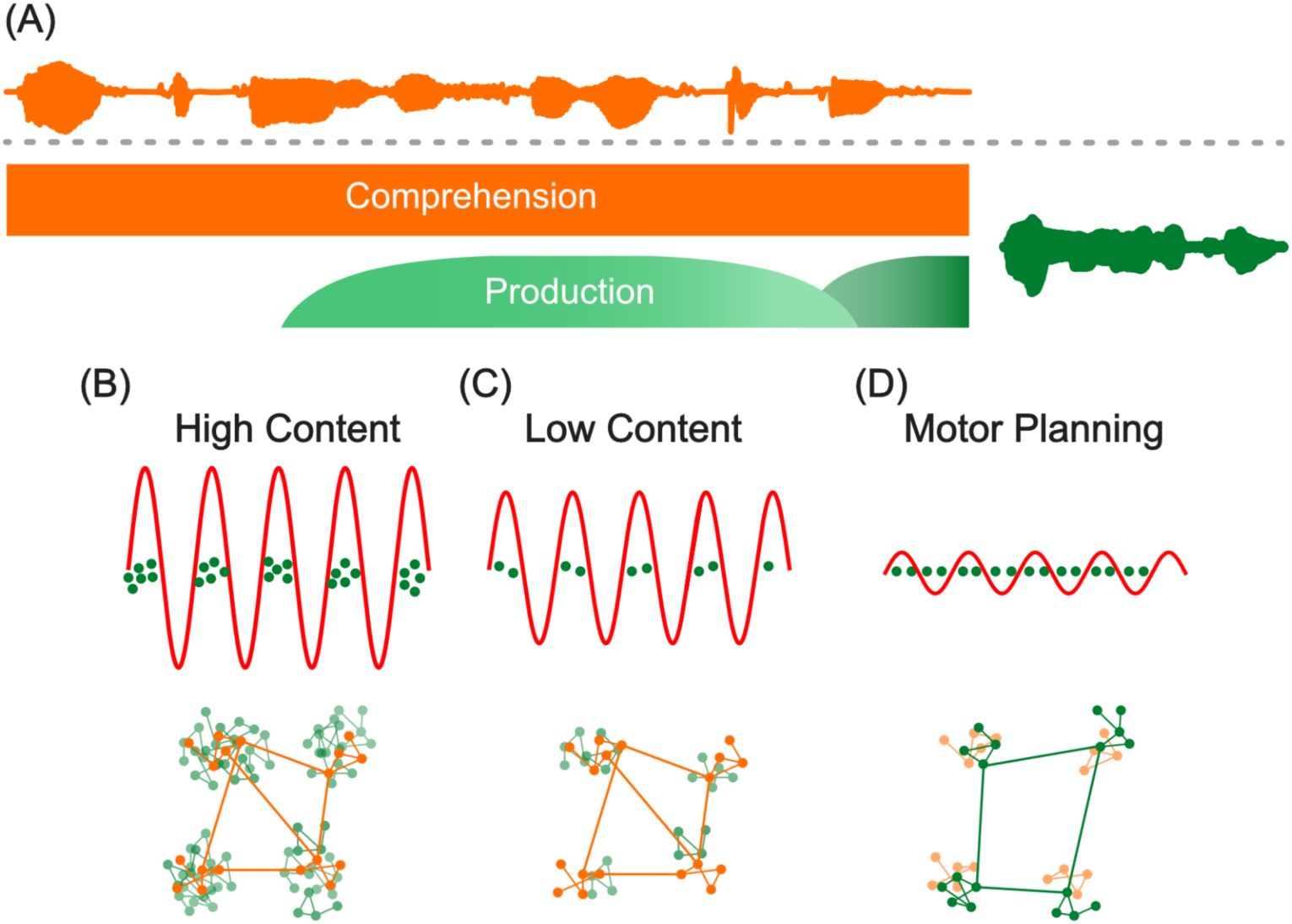
Conceptual model of early maintenance and late commitment during turn preparation. The figure is schematic and intended to illustrate a possible functional organisation consistent with the observed neural dynamics. (A) Speech planning unfolds in two stages: an early content activation stage followed by a later motor-preparation stage. (B) With more content prepared, multiple candidate contents are co-activated and maintained in the background. This is associated with higher alpha power (top) and a larger set of active neural assemblies (bottom). Green assemblies denote production-related processes and orange assemblies denote comprehension-related processes; clusters represent distinct cortical areas. At this stage, only comprehension-related assemblies are bound across areas. (C) With less content prepared, fewer background assemblies are active and alpha power is lower. (D) During late motor preparation, production-related processes become more strongly coordinated as speech onset approaches, while comprehension-related processes recede to the background.

Our findings are broadly compatible with the key assumptions of the *Interactive Alignment Model* (IAM, Pickering and Garrod, 2013). First, IAM assumes some shared resources between comprehension and production mechanisms during dialogue. The presence of behaviourally predictive neural activity during listening is consistent with such overlap. If comprehension and production draw on partially shared representational resources (see also Strijkers and Costa 2016; Pickering and Strijkers 2025), then assembling and maintaining a candidate response while still listening should increase competition within those systems. The sustained pre-speech ERP effects observed here are compatible with this parallel engagement.

Second, IAM proposes that listeners may use production-based mechanisms to anticipate how the partner’s utterance will unfold. Although the present study did not directly manipulate predictability, the early emergence of duration-related activity suggests that response preparation can begin before the partner has finished speaking. This temporal overlap is consistent with the idea that forward modelling or simulation processes can support early preparation in natural conversation, but the present data do not distinguish between predictive simulation and other forms of parallel activation.

IAM further distinguishes between predicting what a partner will say and predicting when their turn will end (Garrod and Pickering 2015). The strongest early neural effects in the present study track the extent of the upcoming response rather than its precise timing. Latency-related predictors were present but more temporally constrained, and partly aligned with partner offset timing. Establishing how timing prediction relates to content prediction will therefore require experimental designs that directly manipulate turn-end cues and contextual information.

### Methodological contribution

Beyond theoretical implications, these results demonstrate that meaningful neural signatures of speech planning can be recovered from EEG recorded during spontaneous, face-to-face conversation. By focussing on the listening periods preceding speech onset, we avoided the most severe speech-related artefacts while preserving ecological validity. Crucially, neural signals during these artefact-minimal intervals predicted turn-level behaviour at the level of individual trials. This shows that natural conversation can yield interpretable, behaviourally grounded neural data, not only coarse measures of interbrain coupling

### Limitations

Although we analysed listening periods preceding speech onset to minimise artefacts, subtle muscle activity near speech onset cannot be fully excluded. Combining EEG with EMG, or using intracerebral recordings, will be valuable for separating cortical activity from peripheral sources.

The conversational nature of the task also increases variability across turns and participants. While we mitigated this by analysing large numbers of turns and using mixed-effects models with behavioural covariates, richer annotation of discourse function and informational content will be important to refine interpretation. Also, because response duration likely covaries with processing demands and complexity, we cannot fully dissociate planning-related activity from general cognitive load or effort.

Finally, Diapix elicits relatively structured collaborative dialogues. Extending the approach to more spontaneous settings will be important to assess generalisability

## Conclusion

In summary, neural activity recorded during spontaneous conversation predicts properties of the next speaking turn, most robustly the duration of the upcoming contribution. Pre-speech ERPs and oscillatory dynamics carry information about response duration more than one second before speech articulation, during the listening period. This extends evidence from controlled laboratory paradigms to natural interaction and supports a view in which response-related representations can be assembled and maintained during comprehension, followed by a later transition toward commitment and execution as onset approaches. More broadly, the results indicate that turn taking is supported by temporally structured neural dynamics that manage competition between listening and speaking and coordinate their interaction in real time.

## Materials and Methods

### Participants

Thirty-four healthy adults (seventeen dyads) took part in the study (mean age = 20.75 years, SD = 3.13; 4 left handed, 6 males). All participants were native French speakers with normal or corrected vision and no known history of neurological anomalies. Within each dyad, participants were sex-matched (female-female or male-male) and unfamiliar with each other prior to the experiment. Facial hair was an exclusion criterion to ensure proper electrode-skin contact for EMG recordings. One participant was excluded due to poor EEG cap placement and one due to EEG recording failure, leaving 32 participants in total. Due to acquisition errors, one participant is missing 2 runs and two participants are missing 1 run. Participants received 35 Euros for their participation. All procedures were approved by the Inserm Ethics Committee (approval no. 2021-A02248-33). Participants received written study information by email and provided informed consent prior to participation, documented by email confirmation, in accordance with the Declaration of Helsinki.

### Task

In order to elicit spontaneous conversational turn taking, we used the Diapix task, a collaborative ‘spot-the-difference’ game originally developed by Van Engen et al. (2010) and refined by Baker and Hazan (2011). Each participant in the dyad is given a similar but not identical image. The task is to find the differences through conversation, without seeing their partner’s image. For each pair of images, the participants were given 4 minutes to find as many differences as possible. This setup naturally encouraged continuous dialogue and frequent turn exchanges.

### Stimuli

The stimuli were adapted from the French version of the Diapix materials (*DiapixFR,* Marmel et al. 2020), supplemented by translation of missing *DiapixUK* images to complete the full set (Baker and Hazan 2011). The pictures depict three scenes (*beach*, *farm* and *street*), with four variants each, yielding 12 image pairs. Each pair of images contained 12 differences, but this was not disclosed to participants to encourage them to continue speaking until the trial ended. For further details of the images see Baker and Hazan (2011).

### Procedure

Upon arrival, both participants were fitted with 64-channel EEG caps. Then they sat face-to-face in the recording room, with electrodes and cables secured to minimise movement related artefacts. Participants were instructed to converse naturally while keeping head and body movements to a minimum. The experiment began with a 1-min eyes-open resting-state recording, followed by eight Diapix conversations (4 min each, pseudo-randomly chosen from the 12 image pairs), and concluded by another 1-min resting state. The beginning and the end of each section was marked by a brief auditory beep. The whole experiment lasted roughly 90 minutes.

### Data Acquisition

#### Audio Acquisition

Both participants were fitted with AKG C520 head-worn condenser microphones on top of the EEG caps. Audio was recorded with Audacity (Version 3.1.3; Audacity Team, 2021) at 48 kHz in 16 bits as a stereo channel with each speaker’s voice in each channel. Data were acquired continually then cropped manually after the experiment.

#### EEG Acquisition

EEG and audio data were recorded simultaneously from both participants during all conversation blocks. EEG was acquired using two BioSemi Active Two 64-channel systems (standard 10/20 layout, BioSemi, the Netherlands) with amplifiers daisy-chained for synchronization. Signals were amplified (−3 dB at ∼204 Hz low-pass, DC coupled) and digitised at 2048 Hz.

#### Speech Transcription

Audio files were first cropped to the 4-min task duration and transcribed with an automatic transcription pipeline (Yamasaki et al. 2023). The resulting transcriptions were manually corrected using Praat 6.4.12 (Boersma and Weenink 2021) then time-aligned at word and phoneme levels using Julius (Lee et al. 2001) and SPPAS (Bigi 2015). The resulting TextGrid annotation provided both word and phoneme level boundaries for each participant.

From these annotations we defined Inter-Pausal Units (IPUs) as speech segments produced by a single speaker bounded by silences of at least 200 ms, excluding non ‘speech’ sounds (e.g. laughter, coughs). This threshold is used to segment continuous speech within a speaker and does not constrain turn-taking events between speakers. We define turn taking as the onset of an IPU by one speaker following the offset of their partner’s IPU, with response latencies between -1000 ms and +1000 ms (see Fig. 3). This time window corresponds to typical floor-transition times in natural conversation (Levinson and Torreira 2015).

#### EEG Preprocessing

EEG data were analysed with MNE-python (Gramfort et al. 2013). Signals were re-referenced to the average of all channels to minimize contamination from speech-related artefacts observable in earlobe channels.

To attenuate muscle and movement artefacts, we applied independent component analysis (ICA) after downsampling data to 512 Hz, high-pass filtering at 1 Hz, and concatenating all conversational blocks. The number of components was set to the number of channels minus identified bad channels. Artefact components were detected automatically using two complementary algorithms: (1) MNE-python’s built-in ‘‘find_bads_musclè’ function classifying muscle related components using spectral slope, peripherality and spatial smoothness (Dharmaprani et al. 2016), and (2) *ICLabel,* a machine-learning classifier trained on a large amount of data to identify a variety of artefacts (Pion-Tonachini et al. 2019; Li et al. 2022). A component flagged by *either method was removed* (mean 32.1, SD = 6.0). Finally, bad channels were interpolated and the final clean signal was band-pass filtered between 0.1 and 30 Hz.

#### EEG data analyses

In the analyses below, *self* refers to the EEG-recorded participant for a given trial, even though the analysed window always corresponds to the period before their own speech onset, that is while they are listening to their partner. Their partner is referred to as the *other-participant*. Each participant serves as *self* in their own listening periods, so all analyses were performed symmetrically for both members of the dyad. All turn-level analyses (ERPs, time-frequency, and mixed-effects models) were based on the same set of 12,939 trials, with per-participant trial counts ranging from 99 to 314.

#### Epoching and event definitions

EEG epochs were time-locked to the onset of the inter-pausal unit (IPU) of the self-participant and spanned from −2200 ms to +500 ms relative to self speech onset (0 ms). IPUs not corresponding to turn taking, or shorter than 10 ms, were excluded.

#### Event-related potential analysis

ERPs were linearly detrended and baseline-corrected using the full epoch window (−2200 to +500 ms), avoiding onset-centred baselines that could introduce condition-dependent bias.

To obtain an initial spatiotemporal map of ERP differences and define analysis windows, trials were split by (1) median response duration (long versus short) and (2) median response latency (fast versus slow). Statistical differences between conditions were assessed using a two-tailed cluster-based spatiotemporal paired t-test implemented in MNE-Python’s ‘‘spatio_temporal_cluster_1samp_test()’’ (Maris and Oostenveld, 2007). The cluster-forming threshold was set to p < 0.05, with 1000 permutations. The spatial adjacency matrix was computed from the standard 10-20 compatible BioSemi 64-channel montage.

#### Time-frequency analyses

Using the same epochs, alpha (8-12 Hz) and beta (13-30 Hz) band-limited activity were estimated by band-pass filtering and computing the amplitude envelope of the analytic signal using the Hilbert transform which was then squared to obtain power. To reduce edge effects, the first 200 ms and the last 500 ms of each epoch were removed. The same cluster-based permutation procedure as for the ERP analyses was used to assess power differences between conditions.

#### Trial-level mixed-effects modelling

To account for individual variability and behavioural covariation, trial-level EEG and behavioural data were entered into linear mixed-effects models (LMMs; lme4, v1.7.2; Bates et al., 2015) in R (v4.4.3). Two dependent variables were analysed separately: response duration (the length of the speaker’s IPU) and response latency (the gap between the partner’s offset and the speaker’s onset). All LMM results are FDR corrected with Benjamini-Hochberg procedure.

##### Behavioural dependency models

To quantify inter-variable dependencies, three behavioural models were fitted:

● self-duration ∼ latency + (1 | participant)
● self-duration ∼ other-duration + (1 | participant)
● latency ∼ other-duration + (1 | participant)

##### Neural predictors and model specification

EEG epochs were time-locked to the onset of the self-participant’s IPU and spanned from −2200 ms to +500 ms relative to speech onset (0 ms). The extended window allowed a 200 ms margin at the beginning of the epoch to minimise edge effects for time-frequency analyses. For statistical analyses, all tests were restricted to the listening interval from −2000 to 0 ms, excluding post-onset activity prone to speech artefacts and avoiding edge-related distortions at the beginning of the epoch.

For time-frequency analyses, alpha- and beta-band power envelopes were estimated from the epoched EEG by band-pass filtering and computing square of the magnitude of the analytic signal using the Hilbert transform. To further reduce edge effects introduced by filtering and Hilbert transformation, the first 200 ms and the last 500 ms of each epoch were excluded prior to power estimation.

Neural predictors were defined as the mean ERP amplitude or oscillatory power within two pre-speech time windows and two regions of interest. These ROIs and time windows were not defined a priori, but were derived from the spatio-temporal clusters identified in the preceding cluster-based permutation analyses.

Specifically, clusters showing significant condition differences were used to guide the selection of anterior and posterior electrode groups and the division of the pre-speech interval into two broad time windows (TW1: −1400 to −800 ms; TW2: −800 to 0 ms).

This approach allowed us to reduce the dimensionality of the data for trial-level mixed-effects modelling while maintaining sensitivity to the main effects revealed by the data-driven cluster analyses.

Behavioural covariates comprised the partner’s previous utterance duration (other-duration) and, for cross-prediction control, the reciprocal behavioural variable (latency when predicting duration, and duration when predicting latency). The resulting model form for duration was:

- self-duration ∼ latency + other-duration + EEG(TW, ROI) + (1 | participant)

The corresponding model form for latency replaced latency with duration as the reciprocal covariate:

- latency ∼ self-duration + other-duration + EEG(TW, ROI) + (1 | participant)

All continuous variables were centred and scaled. Neural models were compared against null models without the neural predictor using likelihood-ratio tests (‘‘anova()’’ in R).

#### ERP-alpha coupling analyses

To examine the relationship between slow ERP activity and alpha power, we conducted additional trial-level analyses focusing on the posterior ROI. First, we tested whether posterior alpha power could be predicted from posterior ERP amplitude using a linear mixed-effects model of the form:

- alpha ∼ ERP + latency + other-duration + (1 | participant) This model was compared to a null model excluding ERP.

Second, to test whether alpha explained variance in response duration beyond ERP amplitude, we compared the following models:

Null model:

self-duration ~ ERP + latency + other-duration + (1 | participant)

Full model:

self-duration ~ ERP + alpha + latency + other-duration + (1 | participant)

Model comparison was performed using likelihood-ratio tests.

#### Decoding and temporal generalisation

To assess the temporal dynamics of neural activity associated with the behavioural variables, we performed temporal generalisation analysis (King and Dehaene, 2014). In temporal generalisation analysis, a model is trained on neural signals at a given time point to classify the target (behavioural) variable, and then tested on neural patterns at all time points, allowing assessment of persisting predictive patterns across time.

EEG epochs were downsampled to 64 Hz, then classified at each time point with a linear support vector machine preceded by per-channel z-scoring across trials (scikit-learn StandardScaler, fit on each training fold only) using 10-fold stratified cross-validation (cross_val_multiscore(), MNE-Python). and evaluated using area under the receiver operating characteristic curve (ROC AUC). Statistical significance was estimated using a one-tailed cluster-based permutation paired t-test against the null hypothesis of AUC = 0.5, using 1000 permutations and p < 0.05 as the cluster-forming threshold.

#### Software

EEG preprocessing and univariate analyses were implemented in MNE-Python (Gramfort et al., 2013). Mixed-effects models were fitted in R (v4.4.3) using lme4 (v1.7.2; Bates et al., 2015). Decoding used scikit-learn via MNE-Python’s cross-validation and temporal generalisation utilities. All analyses were executed via a Snakemake workflow to ensure reproducibility and transparent tracking of dependencies (Köster and Rahmann, 2012).

## Data and Code Availability

The neuroimaging data supporting this study will be made publicly available on OpenNeuro upon publication of the associated data descriptor in Scientific Data. Until then, the data are not publicly available while dataset curation, validation, and repository deposition are ongoing. Additional information is available from the corresponding author upon reasonable request.

Code used for analyses reported in this study is hosted here: https://github.com/CogSci-hiro/turn-taking

## Acknowledgments

Acknowledgments: We thank the participants for their commitment, team members for their help with transcription correction, Sophie Chen for help in experimental setup and data acquisition, Emma Berthault and Anne Mathieu for help with data acquisition and Benjamin Morillon and Kristof Strijkers for comments on the Ms. Research was supported by grants ANR-21-CE28-0010 (DS), ANR16-CONV-0002 (ILCB), ANR-17-EURE-0029 (NeuroMarseille), and the Excellence Initiative of Aix-Marseille University (A*MIDEX).

## Supplementary Materials

### Supplementary Figures

**S1.**
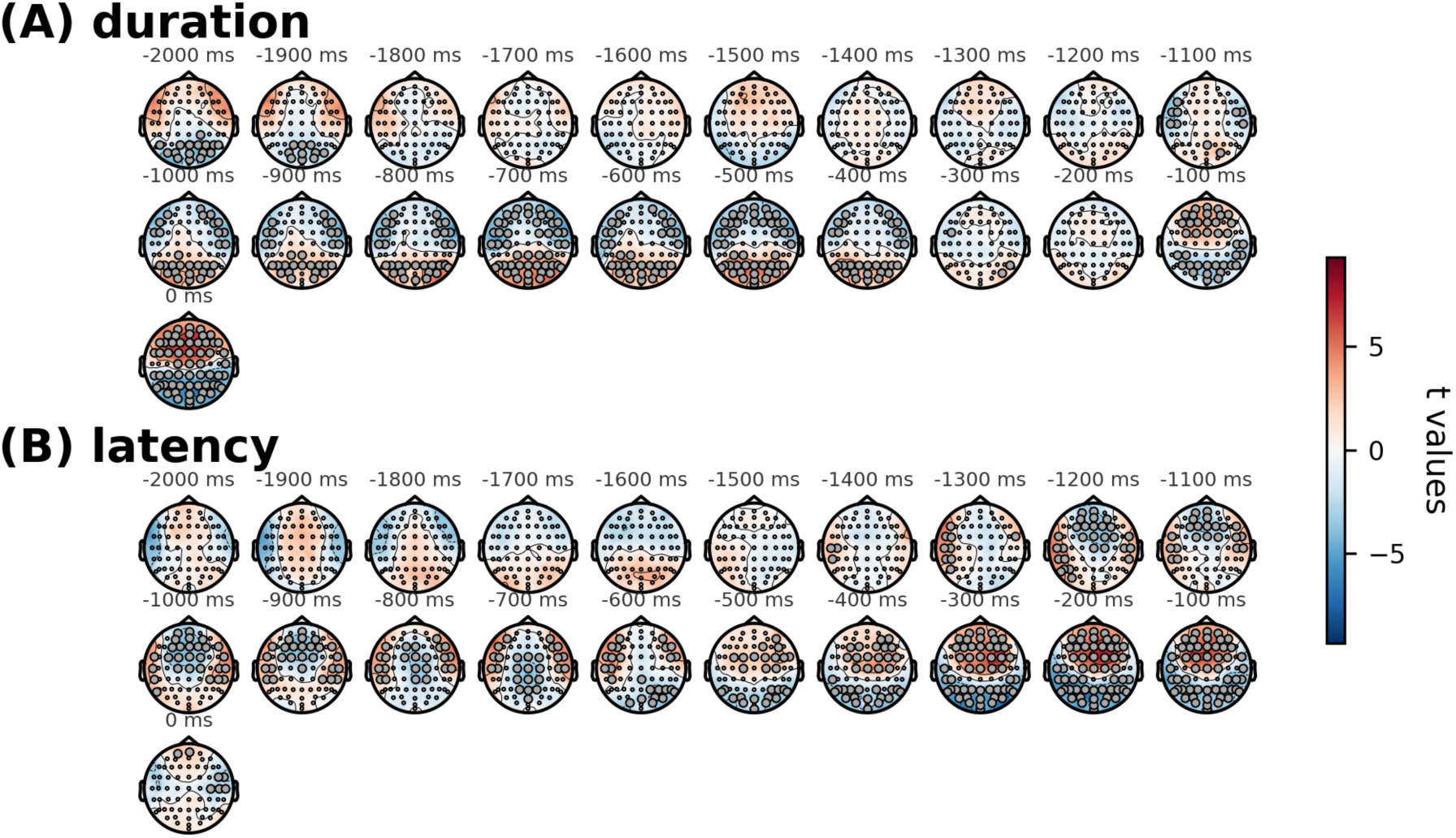
Spatiotemporal distribution of ERP effects for duration and latency conditions. Topographic maps show the scalp distribution of ERP t-values computed for the contrasts of interest. Maps are displayed in 100 ms time steps relative to the event of interest. Panel (A) shows the effect of response duration, and Panel (B) shows the effect of response latency. Warmer colours indicate more positive t-values and cooler colours indicate more negative t-values. Each map represents the spatial distribution of the statistical effect across electrodes within the corresponding time window.

**S2.**
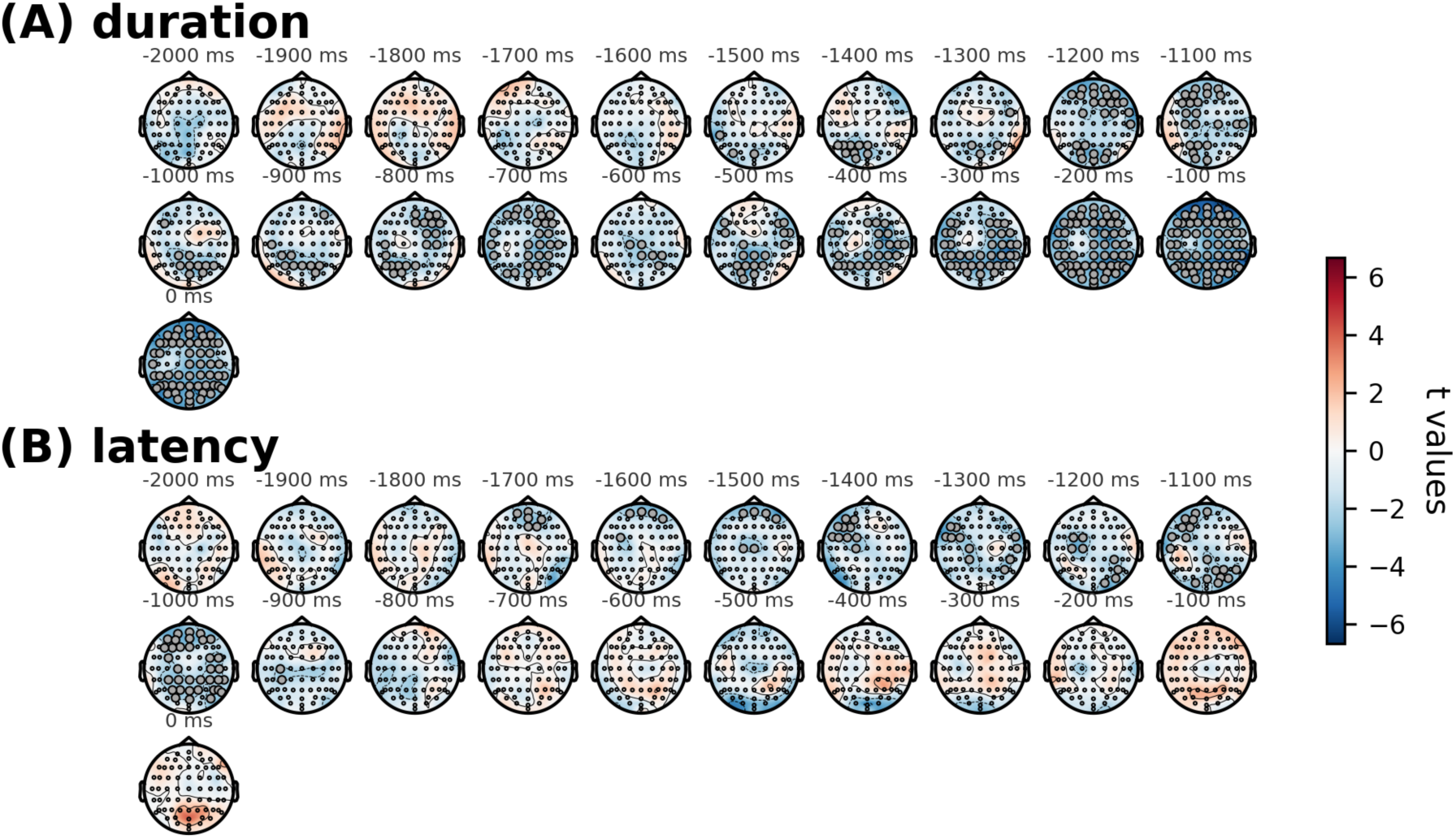
Spatiotemporal distribution of alpha power effects for duration and latency conditions. Topographic maps show the scalp distribution of t-values for alpha-band power computed for the contrasts of interest. Maps are displayed in 100 ms time steps relative to the event of interest. Panel (A) shows the effect of stimulus duration, and Panel (B) shows the effect of response latency. Warmer colors indicate more positive t-values and cooler colors indicate more negative t-values. Each map represents the spatial distribution of the statistical effect across electrodes within the corresponding time window.

**S3.**
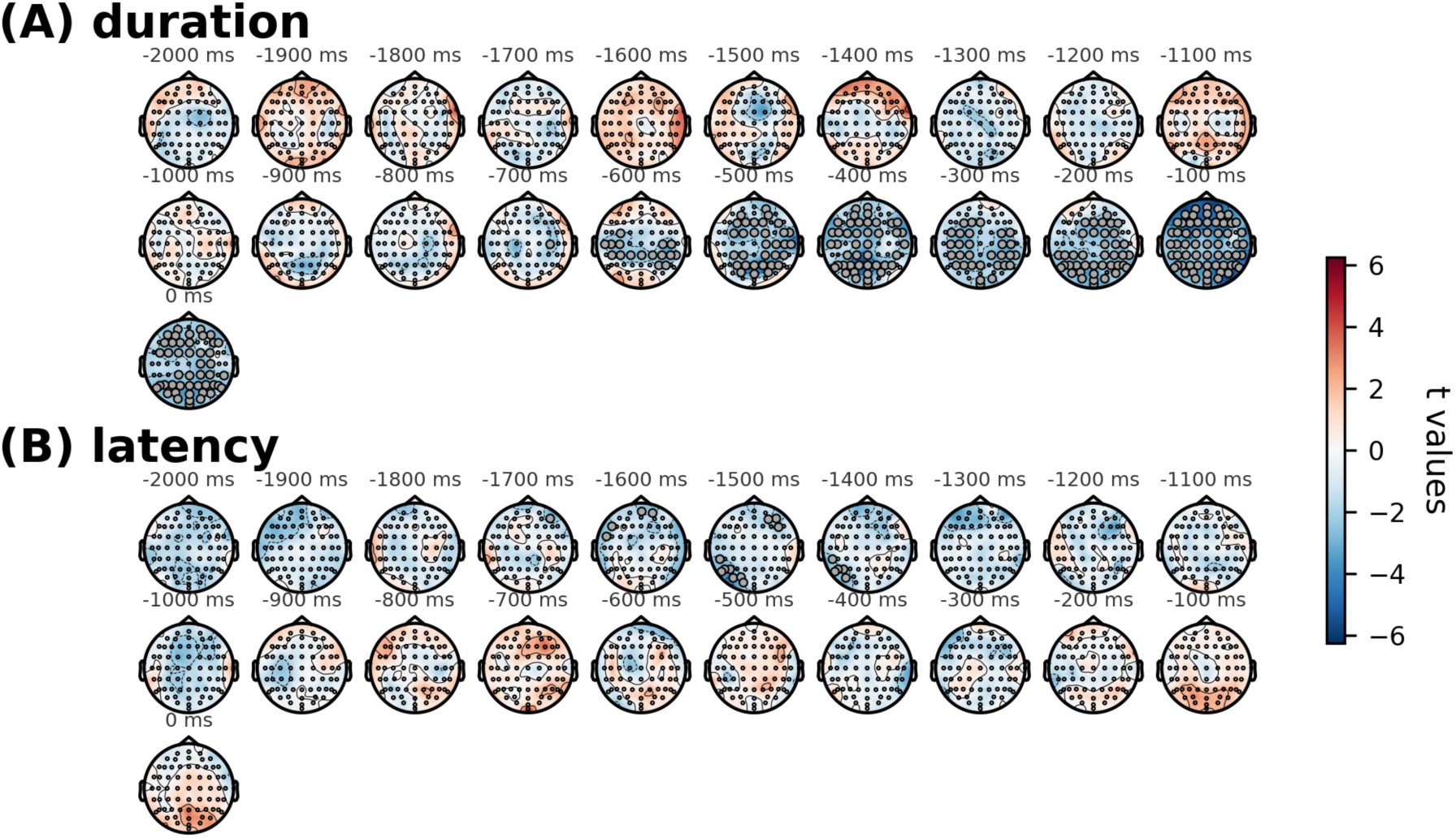
Spatiotemporal distribution of beta power effects for duration and latency conditions. Topographic maps show the scalp distribution of t-values for beta-band power computed for the contrasts of interest. Maps are displayed in 100 ms time steps relative to the event of interest. Panel (A) shows the effect of stimulus duration, and Panel (B) shows the effect of response latency. Warmer colors indicate more positive t-values and cooler colors indicate more negative t-values. Each map represents the spatial distribution of the statistical effect across electrodes within the corresponding time window.

**S4.**
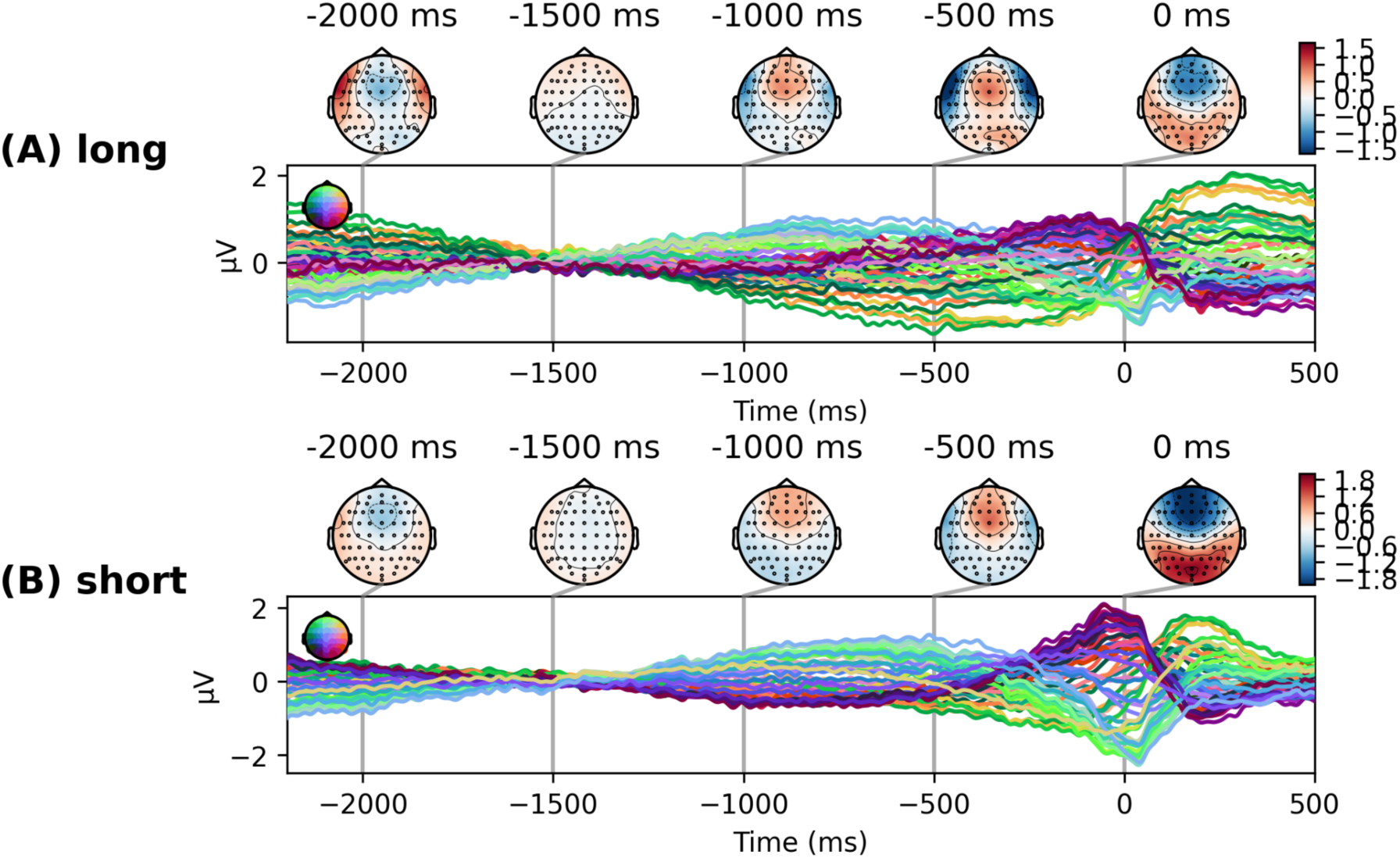
Joint ERP plots by speech duration condition. Joint plots showing the spatiotemporal dynamics of the event-related potential (ERP) for the two speech-duration conditions. (A) Long-duration condition. (B) Short-duration condition. In each panel, the ERP time course is displayed together with scalp topographies at representative latencies, illustrating the spatial distribution of the evoked activity associated with each duration condition.

**S5.**
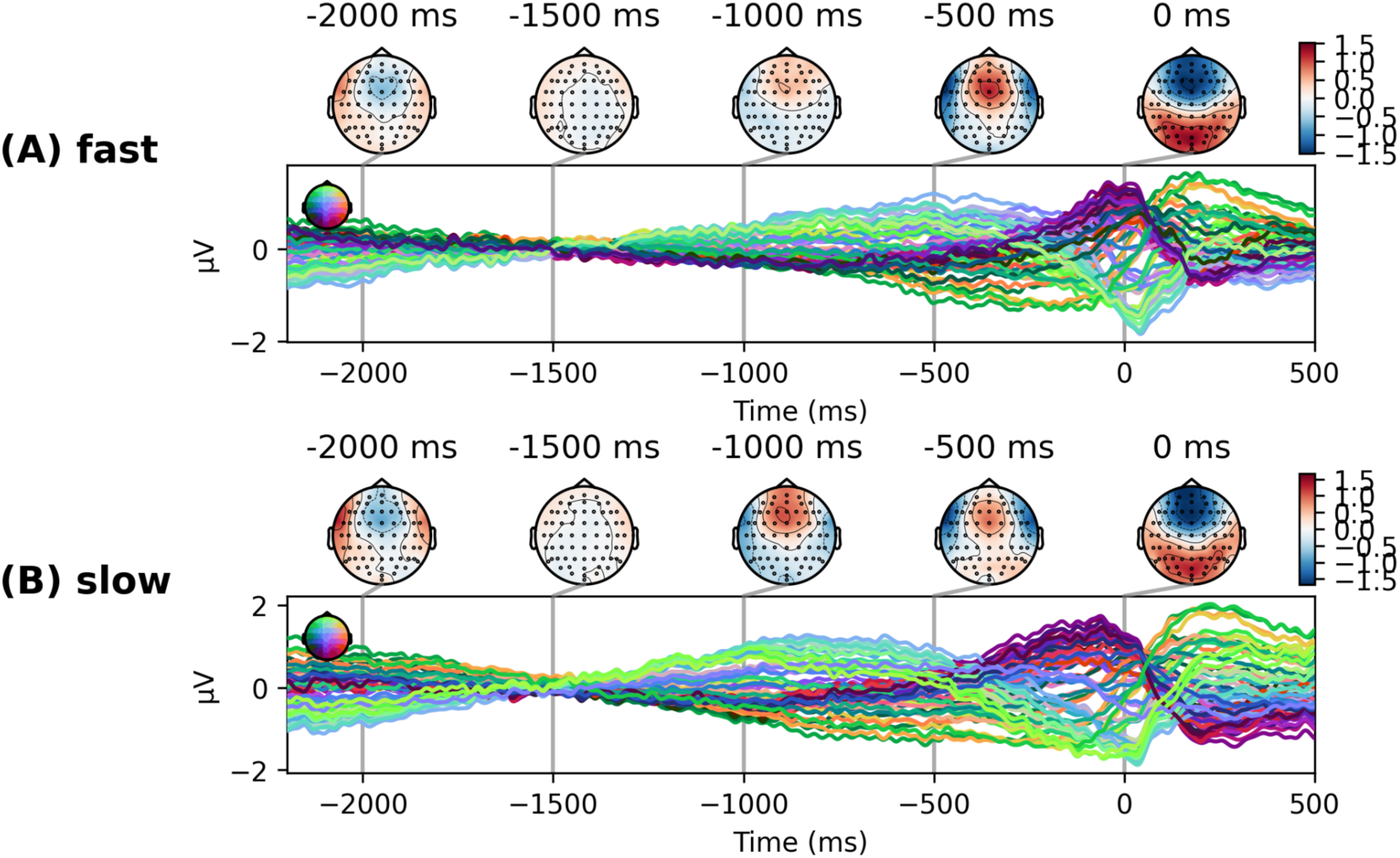
Joint ERP plots by response latency condition. Joint plots showing the spatiotemporal dynamics of the event-related potential (ERP) for the two response-latency conditions. (A) Fast response condition. (B) Slow response condition. In each panel, the time course of the ERP is shown together with scalp topographies at representative latencies, illustrating the spatial distribution of the evoked activity underlying the waveform for each condition.

**S6.**
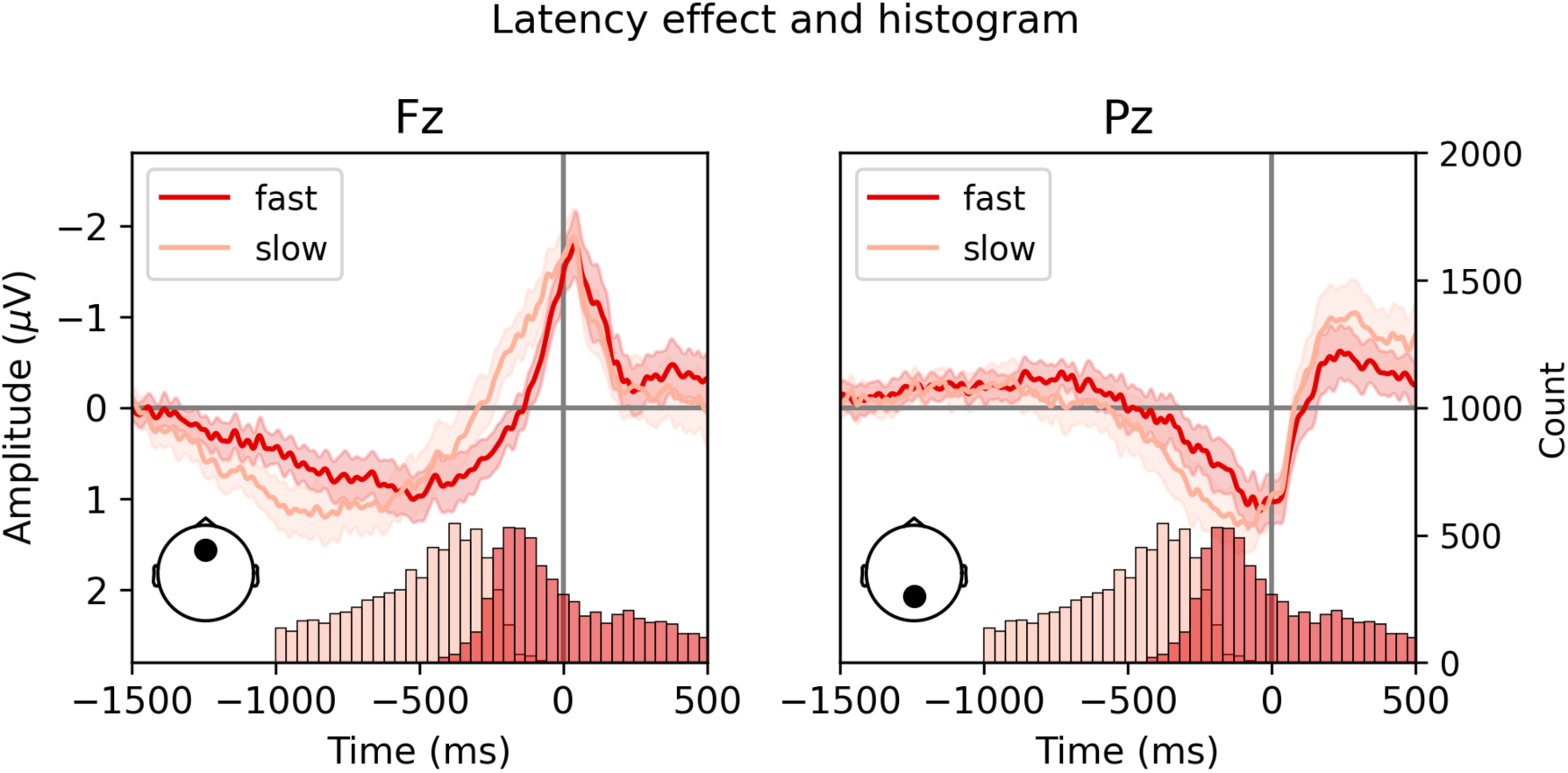
Relationship between ERP morphology and response latency distribution at frontal and parietal electrodes. Event-related potentials (ERPs) were computed after splitting trials by median response latency. Dark red traces indicate trials with faster responses, and light red traces indicate trials with slower responses; shaded areas represent the standard error of the mean. ERPs are shown for electrodes Fz (left) and Pz (right). Histograms below each ERP panel display the distribution of response latencies for the same trials used in the median split. At Fz, the temporal profile of the ERP closely mirrors the latency distribution: the main inflection in the ERP occurs near the peak of the response-latency histogram. This correspondence suggests that the apparent ERP modulation largely reflects alignment to auditory offset timing rather than a genuine latency-related neural difference. In contrast, at Pz the ERP waveform does not follow the latency distribution as closely, indicating that the observed ERP differences more likely reflect a true response-latency-related neural effect rather than a simple consequence of the underlying latency distribution.

